# A Split-Belt Instrumented Treadmill with Uneven Terrain

**DOI:** 10.1101/2024.06.28.601238

**Authors:** Seyed Saleh Hosseini-Yazdi, Arthur D. Kuo

**Affiliations:** Department of Biomedical Engineering, University of Calgary, AB, T2N 1N4 Canada; Faculty of Kinesiology, University of Calgary, AB, T2N 1N4 Canada

**Author notes:** **Correspondence Address:** Dr. Arthur Kuo, Faculty of Kinesiology, University of Calgary, 2500 University Dr NW, Calgary AB T2N1N4, Canada.

## Abstract

The biomechanics of walking are far less understood for uneven terrain than flat or even surfaces. This is due in part to a lack of ground reaction force and moment recordings from each leg. These are often obtained with split-belt instrumented treadmills, which are currently incompatible with uneven terrain, making it difficult to perform biomechanics analyses such as inverse dynamics. Here we show how a standard split-belt instrumented treadmill (Bertec, Inc., Columbus, OH) can be modified to accommodate a variety of uneven terrains. The principal design considerations are structural clearance to allow passage of an uneven treadmill belt and fabrication of the terrain. We designed mechanical components with sufficient clearance for terrains up to 0.045 m high, and formed the terrain from uneven strips of polystyrene. Measured ground reaction forces from each leg at typical walking speeds agreed well with an intact benchmark treadmill (minimum interclass cross correlation score = 0.97). The modifications had negligible effect on the treadmill’s structural strength. The terrain produced some noise-like vibrations, but at much higher frequencies than fundamental to human locomotion. The uneven terrain treadmill can record many steps of the full complement of ground reaction forces and moments from individual legs.

## Introduction

Humans navigate various surfaces in their daily activities including rugged outdoor landscapes. Such surfaces place increased mechanical and energetic demands on the body (Voloshina et al., 2013), but most biomechanical research has focused on flat surfaces, using equipment like force plates and instrumented treadmills (Figure 1 A & B). Treadmills are crucial for measuring ground reaction forces over many walking steps, offering insights into mechanical forces, moments, and powers of the joints (Kuo et al., 2005; Winter, 2009). If an instrumented treadmill could be outfitted with uneven terrain surfaces, new biomechanical measurements could potentially expand the understanding of walking on challenging terrain.

**Figure 1:**
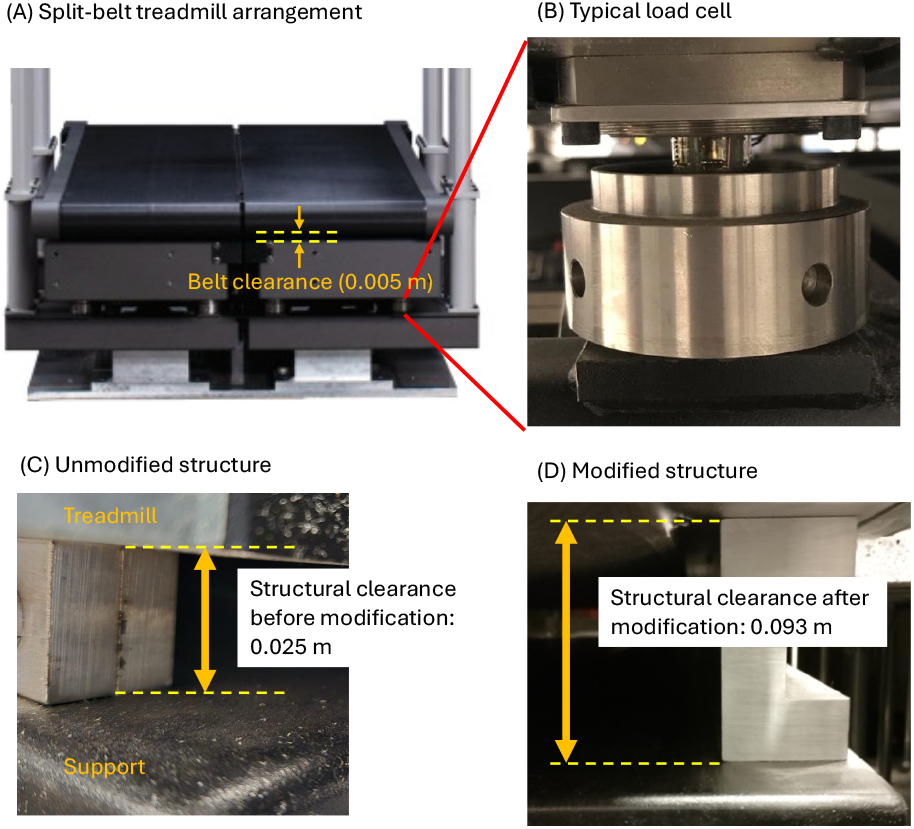
(A) Typical split belt instrumented treadmill arrangement (Bertec Inc., Colombus, OH, USA), with limited structural clearance between the treadmill belt and structure beneath. (B) Each belt is supported by four load cells at four corners. (C) The gap between treadmill conveyor and the supporting structure is small (about 0.025 m). Combined with factors such as roller diameter, the effective clearance between belt and structure is about 0.005 m. (D) A treadmill modification vertically extends the clearance several-fold (to about 0.093 m), allowing for attachment of uneven terrain.

Ground reaction forces have been recorded from uneven walking in limited ways thus far. Voloshina et al. (2013) added uneven terrain to a treadmill placed atop force plates, to record the combined forces of the legs during treadmill walking. They also laid the same treadmill belt atop overground force plates to record individual leg forces, limited however to two steps of overground walking. But uneven walking is less steady than on flat surfaces and is thus best characterized using many steps of data, as with a split-belt instrumented treadmill. Another option involves using shoe insole sensors to estimate individual leg ground reaction forces and pressures in the vertical direction (Downey et al., 2022). However, there is currently no method to record the full complement of individual leg forces and moments during uneven walking with a split-belt instrumented treadmill, the preferred apparatus in many laboratories.

Most instrumented treadmills are not configured for uneven terrain. The terrain used by Voloshina et al. (2013) necessitates extra spatial clearance between the treadmill belt and its supporting structure, to allow unobstructed movement of the uneven bumps around the treadmill’s rollers and underside. However, a typical instrumented treadmill offers very limited belt clearance, for example 0.005 m for a Bertec Instrumented Treadmill (Bertec Inc., Columbus, OH, USA), an amount insufficient for uneven surfaces (Figure 1C).

A split-belt instrumented treadmill adapted to incorporate uneven terrain (Figure 1D) would facilitate controlled biomechanical testing on multiple terrains with varying height differences. We therefore aimed to modify a standard instrumented treadmill (Bertec Inc., Columbus, OH, USA) for greater clearance, to allow fixation of uneven terrain to the belts.

## Methods

We modified an otherwise standard instrumented treadmill for uneven terrain and tested its vibrational characteristics and ability to record individual leg forces. We performed two primary modifications, first a set of structural adjustments to enhance vertical clearance. Second was an improved method to fabricate various uneven surfaces and attach them to the treadmill belt. The structural modifications are specific to an original Bertec instrumented treadmill, but should be adaptable to newer models or other manufacturers. The terrain fabrication is quite general and should be broadly applicable to many treadmills. This section describes the modifications and their associated specifications and computations.

### Structural Modifications

Several modifications were made to increase spatial clearance around the treadmill belt (Figure 2). Typical split-belt instrumented treadmills comprise two self-contained treadmills Figure 2A), each measuring exerted individual leg forces with an independent set of load cells Figure 2B). Most instrumented treadmills have a conveyor atop a supporting structure, usually with limited belt clearance (about 0.005 m for our Bertec treadmill). We increased that clearance by changing the distance between nearest obstructing structures (originally about 0.025 m in Figure 2C, to 0.093 m after modifications, Figure 1D).

**Figure 2:**
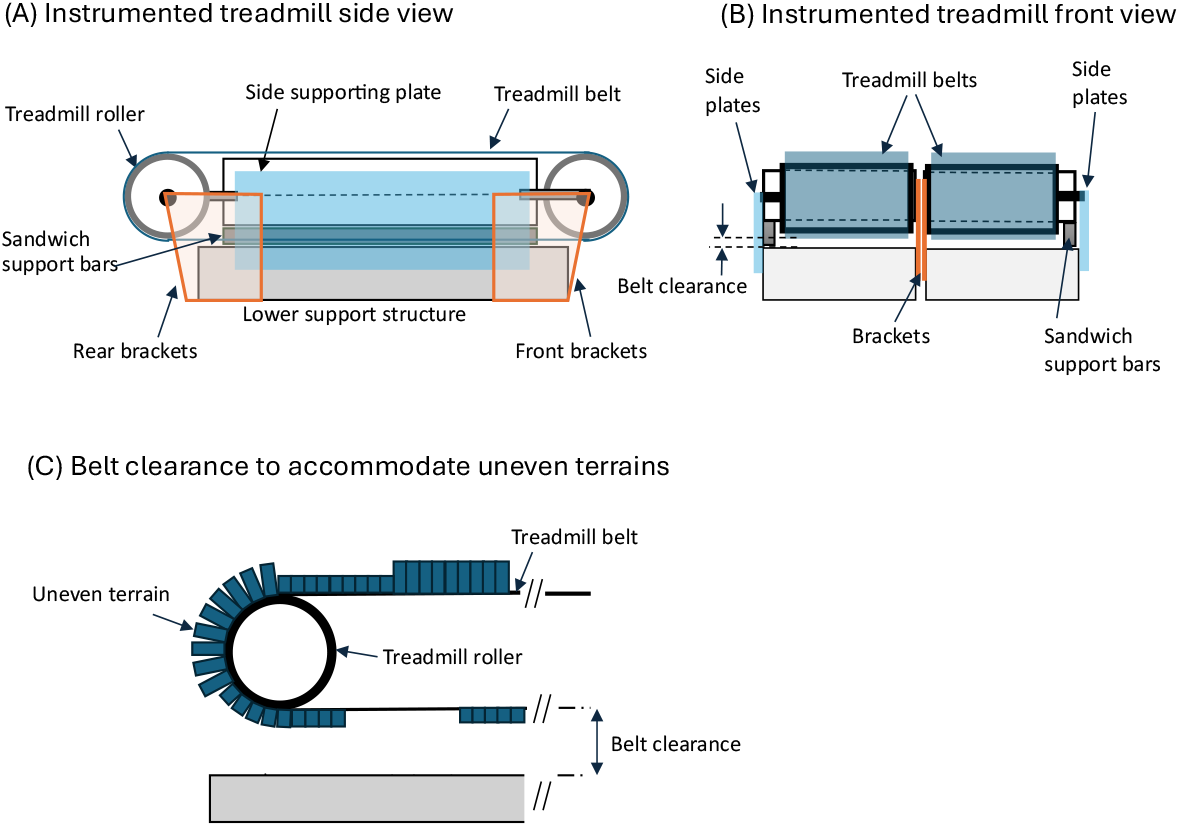
Illustration of instrumented treadmill and need for belt clearance: (A) Unmodified instrumented treadmill side view and (B) front view. (C) To accommodate uneven terrain, additional clearance is needed beneath the conveyor to allow passage of the terrain.

The conveyor frame is connected to a supporting structure below with two narrow supporting assemblies to each side: Sandwich support bars covered by a side plate on the lateral side and two brackets (front and rear) holding the rollers on the medial side. The medial side assembly must be strong and narrow to permit tight medio-lateral clearance between the two treadmill belts. Our design considerations were to improve clearance (Figure 2C) while maintaining structural strength and compatibility with a pulley-driven drive belt between the motors below and the rollers above.

A total of four parts were retrofitted to increase the vertical clearance (Figure 3A). A structural clearance of 0.093 m was specified to provide clearance for the belt as it passes around and under the conveyor, while maintaining compatibility with standard, commercially available timing (drive) belts. We sought at least 6 cm of additional clearance to make it possible to construct terrain of similar amplitude to previous studies (Darici & Kuo, 2022, 2023). To accomplish this, the side plate (A1060 aluminum alloy) and brackets (A36) were replaced with a similar plate of greater height (A6061 and A36 respectively). The sandwich support bars (A1060) were replaced with a single C channel of the same length but taller height (A6061). Thus, the new parts fabricated for each treadmill side were a side plate, front and rear brackets, and a C channel (Figure 3B). Engineering drawings are provided in a publicly accessible repository (Hosseini-Yazdi & Kuo, 2024).

**Figure 3:**
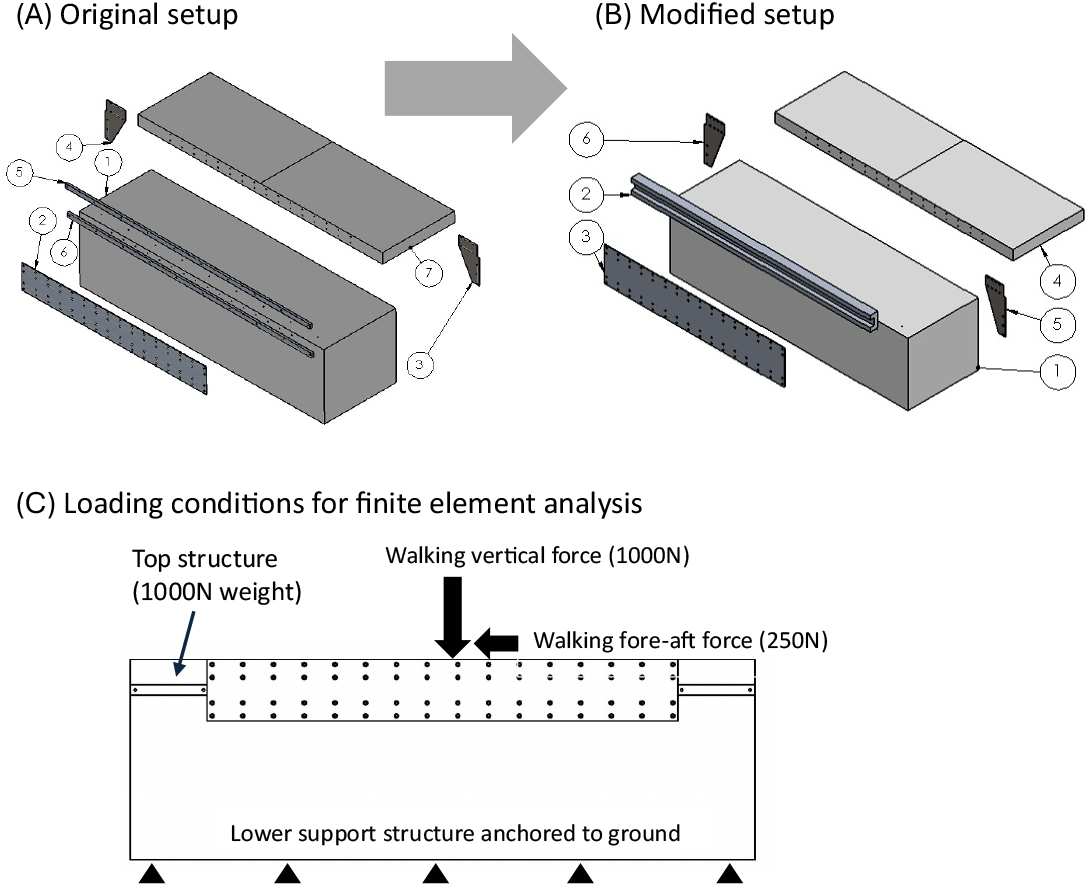
The instrumented treadmill 3D model: (A) the original setup: (A1) supporting bottom structure, (A2) side plate (lateral), (A3) front bracket (medial), (A4) rear bracket (medial), (A5) sandwich support bar (top, lateral), (A6) sandwich support bar (bottom, lateral), (A7) treadmill belt. (B) Modified (vertically extended) belt support: (B1) supporting bottom structure, (B2) C channel to replace the sandwich support bars (lateral), (B3) side plate (lateral), (B4) treadmill belt, (B5) front bracket (medial), (B6) rear bracket (medial). (C) Loading conditions for finite element analysis. Walking load in fore-aft direction: 250 N, in vertical direction: 1000 N, and conveyor weight (distributed dead weight): 1000 N (including a safety factor of 1.5).

We performed Finite Element Analysis (FEA) to assess the strength (static and buckling) and frequency response of the treadmill after the modification (SolidWorks 2020, Dassault Systèmes). The same analysis was conducted on the original structure serving as a benchmark. The analysis loads were representative of a typical walking adult, and consisted of 1000 N in the vertical direction and 250 N in the longitudinal direction to simulate the ground reaction forces experienced during human walking (Collins et al., 2009; Kirtley, 2006). Additionally, we assumed the weight of the treadmill’s belt assembly to be 1000 N, factoring in a minimum safety factor of 1.5 for its dead weight (Figure 3C).

The FEA analysis revealed that the new parts could easily provide sufficient strength. For example, the maximum estimated stress in the front and rear brackets was increased negligibly (original 1.26 MPa to new 1.27 MPa), over sixty times the yield stress limit, and associated displacement kept very small (original 1.5e-7 m to new 4.6e-7 m). The first buckling mode was predicted to occur in the brackets, with a safety factor reduced from 1.5e54 to 1.1e5. The first mode shape’s natural frequency was reduced from the original 3478 Hz to 1609 Hz, well above the frequency content of measured forces (see Results). None of these differences were expected to substantially detract from strength or performance, as the structure can withstand many times the typical adult body weight.

### Terrain Fabrication

The uneven terrain was created by attaching uneven polystyrene blocks to custom-made treadmill belts. Our instrumented treadmills are mounted side-by-side, limiting access to remove and replace the belts to the lateral sides and top. We opted for open-ended belts that could be wrapped around the existing belt and then closed at the ends using a commercial belt lacing system (Alligator Lacing Clips, FLEXCO, Downers Grove, IL, USA). The terrain materials were commercially available treadmill belts (width: 0.406 m, thickness: 0.0018 m, surface diamond pattern, material: PVC). The replacement process involved releasing the treadmill’s rear tensioners, wrapping an uneven terrain belt around the original belt, inserting the lacing pin to connect the ends, and finally re-tensioning the belts by extending the tensioner bolts. The replacement procedure could be completed in about 15 min.

The uneven terrain blocks were of three maximum heights (0.019, 0.032, and 0.045 m), constructed from polystyrene foam sheets. The design criteria were sufficient compressive strength to withstand human walking and low mass to avoid high centripetal forces when going around the treadmill rollers. Each block’s total thickness was produced from two layers of polystyrene, the top being a thin (4.8 mm, Pacon foam boards, Dixon Ticonderoga Company, Appleton, Wisconsin, USA) sheet with paper facing for protection against shoes, and the bottom a similar but bare sheet (various thicknesses). The bottom layer provided most of the height and was cut from standard rigid insulation sheets used for building construction (FOAMULAR NGX C-200 XPS Rigid Foam Insulation Board, Owens Corning, Toledo, OH): strength = 140 kPa, compressive modulus: 6895 kPa, density 25 kg/m^3^). Assuming a shoe initial contact area of 0.01 m^2^, a vertical load of 1000 N, and a compressive stress safety factor of 1.4, a typical peak stress of 100 kPa would cause 6.5e-4 m deformation for the maximum profile height of 0.045 m.

The blocks were cut into strips and glued crosswise onto the treadmill belt. The strips were 0.025 m by 0.406 m and arrayed into groups of twelve to make 0.3 m long blocks in the walking direction (Figure 4A). The strips allowed for movement about the rollers (Voloshina et al., 2013; Voloshina & Ferris, 2018). We used construction glue (LePage PL9000, Henkel Corp., Stamford, CT, USA) to affix the layers to each other and to the belt. The block heights were varied along the walking direction, so that each foot could potentially land flat on one block, or not flat across at most two different ones (Figure 4B). The material was found to be brittle rather than compliant (Figure 4C). Four sets of terrain were constructed, each with a designated maximum foam block height of 0 m (no foam), 0.019 m, 0.032 m, and 0.045 m. The terrains weighed up to 1.2 kg per belt and were found to remain well affixed at a treadmill speed of 1.4 m · s^−1^. Extended walking trials revealed that the foam blocks were sturdy enough for the uneven walking experiments. However, if a subject stepped on a strip as it traveled over either treadmill roller, the strips would occasionally detach from the belt and require regluing.

**Figure 4:**
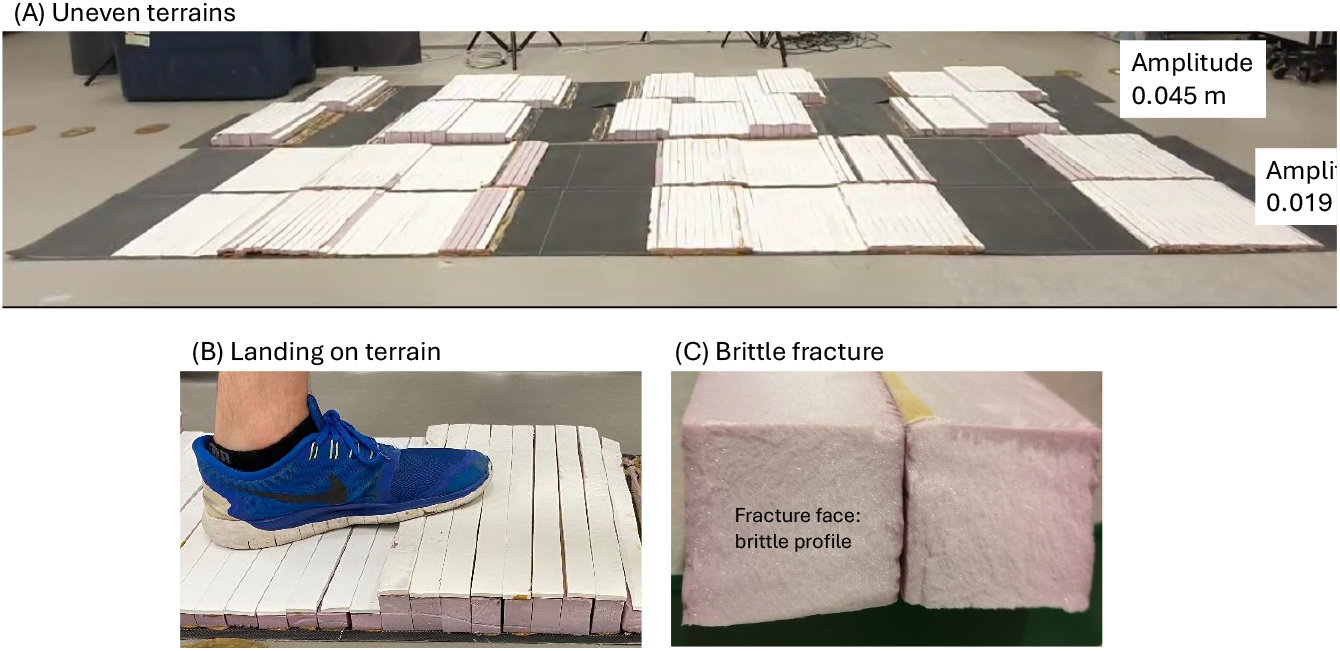
Two examples of uneven terrain belts, as matched pairs laid on the floor for illustration. (A) The terrains were constructed from two layers of extruded polystyrene (white and pink), cut into strips laid mediolaterally atop and glued to each belt. During experiments, the belts were mounted on treadmill with a random relative fore-aft offset, to make the terrain less regular than shown here. (B) A typical landing pattern for the foot. (C) The polystyrene deformed very little under human walking load due to non-compliant, brittle properties. This is illustrated by a brittle fracture face from intentionally broken polystyrene.

## Results of treadmill testing

The modified instrumented treadmill was experimentally tested for mechanical characteristics following structural modification. The tests showed that the uneven terrain belts could withstand walking without slipping, that the treadmill could reproduce the commanded speeds, and that the measured forces were sufficiently accurate. The testing procedure was part of a human subject experiment approved by the University of Calgary Conjoint Health Research Ethics Board, and subjects provided written informed consent before the experiments. Prior to testing, we calibrated the modified treadmill force measurements using a method outlined by Collins et al. (2009).

We found the modifications to have little effect on belt slippage. Slippage was a concern due to the modified belt configuration, where each uneven terrain belt was wrapped around the original bare belt and therefore did not directly contact the drive roller. We first performed a static test by manually grasping and pulling on the belts at up to 359 N as measured by treadmill load cells, which exceeds a typical fore-aft force of about 250 N during walking (Collins et al., 2009). We did not discern any slippage, through visual inspection. Second was a test of treadmill walking where the belt speed was measured and examined for slippage. Young adult subjects (two females and three males, with age 33.6 ± 7.2 yr, weight 70.2 ± 13.5 kg, mean ± s.d.) wore running shoes and walked on the treadmill at five belt speeds (0.6, 0.8, 1.0, 1.2, and 1.4 m · s^−1^). The belt actual traveling speeds were calculated from motion-capture data, with optical markers attached to the belt surfaces (PhaseSpace, San Leandro, CA, USA) at a sampling rate of 960 Hz. We tested all speeds on each terrain, which was wrapped and tensioned around the original smooth belt. We found the measured belt speeds agreed with commanded speeds within about 4%, with coefficient of variation at most about 2%, similar to unmodified. We thus did not find human walking to cause appreciable slipping of uneven terrain belts.

Tested force data also indicate sufficient accuracy of the instrumented force measurements. This was performed because structural modifications could potentially affect load cell accuracy or mechanical vibrations of the entire structure. The modified treadmill’s forces were compared with a similar unmodified treadmill as benchmark, during even walking at 1.2 m · s^−1^ for one subject (Modified and Benchmark, Figure 5; an example of uneven walking forces is also shown). We found the subject’s weight to be within 0.46% between treadmills (modified 73.38 kg, unmodified benchmark 73.04 kg), comparable to reported errors for previous instrumented treadmills (Collins et al., 2009). We also found the force signals from walking to have interclass cross correlation score of at least 0.97.

**Figure 5:**
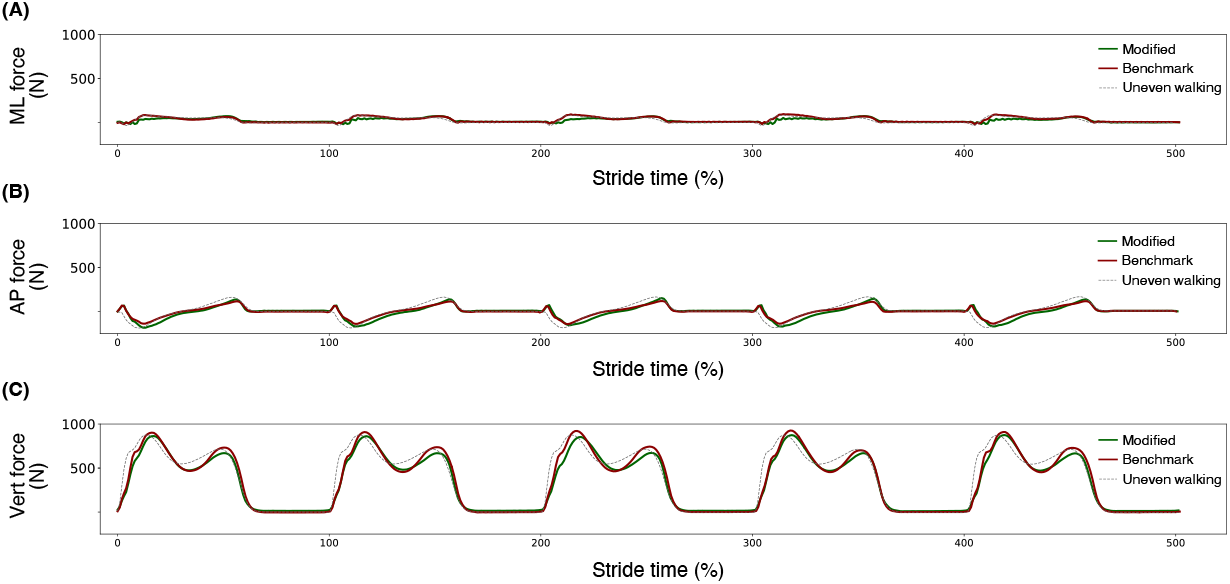
Comparison of ground reaction force components of even walking (1.2 m · s^−1^) for the benchmark and modified treadmills, plus an example of uneven walking. (A) Medio-lateral (ML), (B) antero-posterior (AP), and (C) vertical (Vert) forces. The modified and benchmark signals minimum interclass cross correlation score was 0.97. Additional force traces (light dashed lines) show forces during uneven walking, on synthetic uneven terrain of peak-to-peak amplitude 0.045 m.

We also found the frequency content of collected force data to be acceptable. Power spectral analysis on raw force signals (no filtering) was performed to assess the similarity of force traces (Figure 6). The spectra of the modified and benchmark treadmills appeared similar for a range 0 – 15 Hz typical of walking. Above that range, we observed some downward shifts in vibration frequencies, from 111.9 Hz to 103.4 Hz in the vertical direction, from 64 Hz to 52 Hz in the anteroposterior direction, and from 39 Hz to 33 Hz in the medio/lateral direction. These peaks were 0.52%, 1.08%, and 4.04% of the first fundamental magnitude, respectively. The modified treadmill therefore introduced small, noise-like force fluctuations, but at frequencies at least eighteen times a typical walking stride frequency (1.0 Hz). We associate most of these differences to vibrations and inertial forces of uneven terrain.

**Figure 6:**
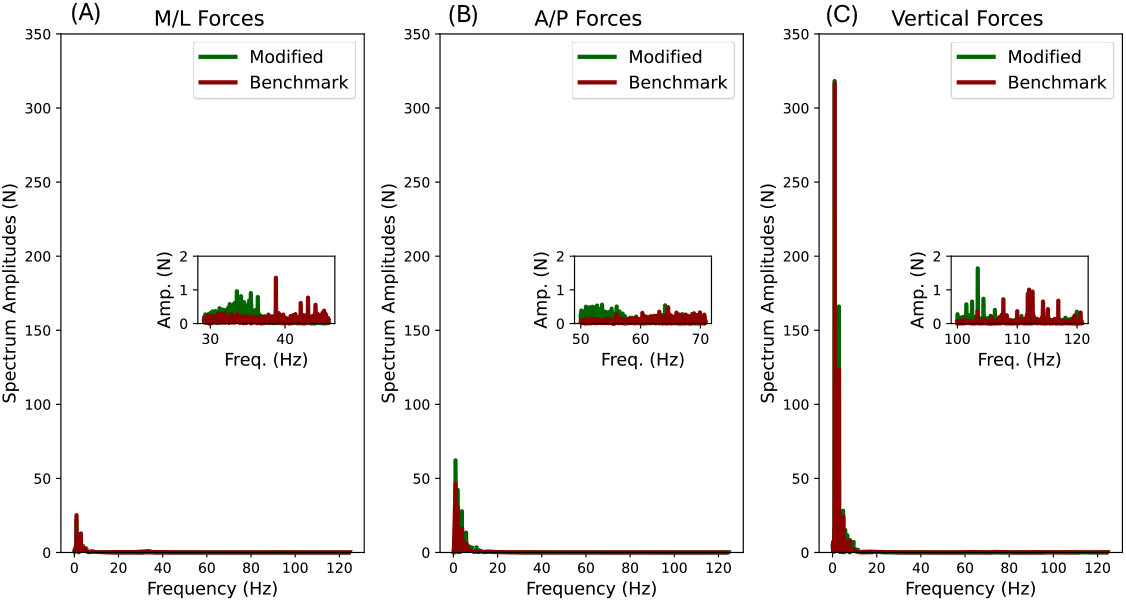
Spectrum analysis of benchmark and modified treadmills’ ground reaction force components for even walking (at 1.2 m · s^−1^). (A) Medio-lateral, (B) antero-posterior, and (C) vertical spectra. The spectra were similar at frequencies below about 15 Hz, about 15 times a typical walking stride frequency of about 1 Hz. The modifications did cause some vibrations to occur at other frequencies, generally above 30 Hz (see inset).

## Discussion

Split-belt instrumented treadmills have enabled measurement of ground reaction forces for analyses such as inverse dynamics. This capability has now been extended to uneven terrain surfaces of varying amplitude, allowing for controlled terrain studies not previously possible. The present device improves on previous solutions and offers new avenues for biomechanical investigation.

Our procedure differs from a previous uneven terrain treadmill (Voloshina et al., 2013) in three main aspects. First is the application to a split-belt treadmill for individual leg forces, rather than a single treadmill measuring only combined forces. A second difference is the use of a reversible lacing system for the belts, to enable switching between different uneven terrains (about 15 min per replacement) within a single experiment. An alternative procedure without belt replacement is to affix individual terrain obstacles to a permanent belt with hook-and-loop fasteners (Downey et al., 2022), which allows for greater flexibility in terrain configuration and shape, but perhaps longer switching time.

The third difference is in the use of polystyrene foam for the uneven terrain. Polystyrene is much lighter than the previous wooden base (sugar pine wood, about sixteen times denser) (Voloshina et al., 2013). It greatly reduces inertial forces as the strips travel around the treadmill rollers that can cause detachment from the belt. We found polystyrene to be sufficiently durable for many hours of experimental trials, with occasional need for re-gluing of detached, broken, or separated strips. Greater terrain amplitudes than those examined here may also be possible but have yet to be tested.

Some care must be taken with this uneven terrain. Subjects should be instructed not to step on the terrain strips at either far end of the treadmill as they travel over a roller, which may cause detachment. Polystyrene is weaker than wood, and so concentrated stresses should also be avoided. The terrain was protected by paper facing on the polystyrene and by instructing participants to wear soft soled shoes. Although polystyrene’s elastic modulus is not high (6895 kPa), the material is inelastic, as evident from a brittle failure mode (Figure 4C). Informal tests suggest that this terrain is also acceptable for running, although faster speeds cause greater deformation and more frequent detachment. Stronger material such as balsa wood and stronger fixing methods than glue may be preferable for extended running studies (Voloshina et al., 2013; Voloshina & Ferris, 2018).

A split-belt instrumented treadmill with uneven terrain makes it possible to collect many steps of force data during walking. Such capability may be especially helpful for evaluating uneven and unsteady walking in populations such as older adults (Downey et al., 2022) and amputees (Paysant et al., 2006; Stewart et al., 2024). We expect the modifications described here to be adaptable to other treadmills and other uneven surface compositions, including compliant surfaces and energy-absorbing ones. There is considerable potential to apply inverse dynamics and force-based analyses, conventionally applied only to even surfaces, to uneven, compliant, or energy-absorbing surfaces.

## Acknowledgements

This work was supported in part by the Natural Sciences and Engineering Research Council of Canada (NSERC) Discovery and Canada Research Chair (Tier 1) programs.

